## Supplementary material for "scARE: Attribution Regularization for Single Cell Representation Learning": LaTeX: example_paper.bib

@article{erion2021improving,
  title={Improving performance of deep learning models with axiomatic attribution priors and expected gradients},
  author={Erion, Gabriel and Janizek, Joseph D and Sturmfels, Pascal and Lundberg, Scott M and Lee, Su-In},
  journal={Nature machine intelligence},
  volume={3},
  number={7},
  pages={620--631},
  year={2021},
  publisher={Nature Publishing Group UK London}
}

@article{lopez2018deep,
  title={Deep generative modeling for single-cell transcriptomics},
  author={Lopez, Romain and Regier, Jeffrey and Cole, Michael B and Jordan, Michael I and Yosef, Nir},
  journal={Nature methods},
  volume={15},
  number={12},
  pages={1053--1058},
  year={2018},
  publisher={Nature Publishing Group US New York}
}

@article{kingma2015variational,
  title={Variational dropout and the local reparameterization trick},
  author={Kingma, Durk P and Salimans, Tim and Welling, Max},
  journal={Advances in neural information processing systems},
  volume={28},
  year={2015}
}

@article{ashuach2022peakvi,
  title={PeakVI: A deep generative model for single-cell chromatin accessibility analysis},
  author={Ashuach, Tal and Reidenbach, Daniel A and Gayoso, Adam and Yosef, Nir},
  journal={Cell reports methods},
  volume={2},
  number={3},
  pages={100182},
  year={2022},
  publisher={Elsevier}
}

@article{gayoso2021joint,
  title={Joint probabilistic modeling of single-cell multi-omic data with totalVI},
  author={Gayoso, Adam and Steier, Zo{\"e} and Lopez, Romain and Regier, Jeffrey and Nazor, Kristopher L and Streets, Aaron and Yosef, Nir},
  journal={Nature methods},
  volume={18},
  number={3},
  pages={272--282},
  year={2021},
  publisher={Nature Publishing Group US New York}
}

@article{heumos2023best,
  title={Best practices for single-cell analysis across modalities},
  author={Heumos, Lukas and Schaar, Anna C and Lance, Christopher and Litinetskaya, Anastasia and Drost, Felix and Zappia, Luke and L{\"u}cken, Malte D and Strobl, Daniel C and Henao, Juan and Curion, Fabiola and others},
  journal={Nature Reviews Genetics},
  pages={1--23},
  year={2023},
  publisher={Nature Publishing Group UK London}
}

@article{xu2021probabilistic,
  title={Probabilistic harmonization and annotation of single-cell transcriptomics data with deep generative models},
  author={Xu, Chenling and Lopez, Romain and Mehlman, Edouard and Regier, Jeffrey and Jordan, Michael I and Yosef, Nir},
  journal={Molecular systems biology},
  volume={17},
  number={1},
  pages={e9620},
  year={2021}
}

@article{sikkema2023integrated,
  title={An integrated cell atlas of the lung in health and disease},
  author={Sikkema, Lisa and Ram{\'\i}rez-Su{\'a}stegui, Ciro and Strobl, Daniel C and Gillett, Tessa E and Zappia, Luke and Madissoon, Elo and Markov, Nikolay S and Zaragosi, Laure-Emmanuelle and Ji, Yuge and Ansari, Meshal and others},
  journal={Nature Medicine},
  pages={1--15},
  year={2023},
  publisher={Nature Publishing Group US New York}
}

@article{hesse2021fast,
  title={Fast axiomatic attribution for neural networks},
  author={Hesse, Robin and Schaub-Meyer, Simone and Roth, Stefan},
  journal={Advances in Neural Information Processing Systems},
  volume={34},
  pages={19513--19524},
  year={2021}
}

@article{buenrostro2015single,
  title={Single-cell chromatin accessibility reveals principles of regulatory variation},
  author={Buenrostro, Jason D and Wu, Beijing and Litzenburger, Ulrike M and Ruff, Dave and Gonzales, Michael L and Snyder, Michael P and Chang, Howard Y and Greenleaf, William J},
  journal={Nature},
  volume={523},
  number={7561},
  pages={486--490},
  year={2015},
  publisher={Nature Publishing Group UK London}
}

@article{stoeckius2017simultaneous,
  title={Simultaneous epitope and transcriptome measurement in single cells},
  author={Stoeckius, Marlon and Hafemeister, Christoph and Stephenson, William and Houck-Loomis, Brian and Chattopadhyay, Pratip K and Swerdlow, Harold and Satija, Rahul and Smibert, Peter},
  journal={Nature methods},
  volume={14},
  number={9},
  pages={865--868},
  year={2017},
  publisher={Nature Publishing Group}
}

@article{de2022population,
  title={Population-level integration of single-cell datasets enables multi-scale analysis across samples},
  author={De Donno, Carlo and Hediyeh-Zadeh, Soroor and Wagenstetter, Marco and Moinfar, Amir Ali and Zappia, Luke and Lotfollahi, Mohammad and Theis, Fabian J},
  journal={bioRxiv},
  pages={2022--11},
  year={2022},
  publisher={Cold Spring Harbor Laboratory}
}

@article{teichmann2020method,
  title={Method of the Year 2019: single-cell multimodal omics},
  author={Teichmann, Sarah and Efremova, Mirjana},
  journal={Nat. Methods},
  volume={17},
  number={1},
  pages={2020},
  year={2020}
}

@article{kuemmerle2022probe,
  title={Probe set selection for targeted spatial transcriptomics},
  author={Kuemmerle, Louis B and Luecken, Malte D and Firsova, Alexandra B and Barros de Andrade e Sousa, Lisa and Stra{\ss}er, Lena and Heumos, Lukas and Mekki, Ilhelm Isra and Mahbubani, Krishnaa T and Sountoulidis, Alexandros and Balassa, Tamas and others},
  journal={bioRxiv},
  pages={2022--08},
  year={2022},
  publisher={Cold Spring Harbor Laboratory}
}

@article{lotfollahi2023biologically,
  title={Biologically informed deep learning to query gene programs in single-cell atlases},
  author={Lotfollahi, Mohammad and Rybakov, Sergei and Hrovatin, Karin and Hediyeh-Zadeh, Soroor and Talavera-L{\'o}pez, Carlos and Misharin, Alexander V and Theis, Fabian J},
  journal={Nature Cell Biology},
  volume={25},
  number={2},
  pages={337--350},
  year={2023},
  publisher={Nature Publishing Group UK London}
}

@inproceedings{qoku2023encoding,
  title={Encoding Domain Knowledge in Multi-view Latent Variable Models: A Bayesian Approach with Structured Sparsity},
  author={Qoku, Arber and Buettner, Florian},
  booktitle={International Conference on Artificial Intelligence and Statistics},
  pages={11545--11562},
  year={2023},
  organization={PMLR}
}

@article{conard2023spectrum,
  title={A spectrum of explainable and interpretable machine learning approaches for genomic studies},
  author={Conard, Ashley Mae and DenAdel, Alan and Crawford, Lorin},
  journal={Wiley Interdisciplinary Reviews: Computational Statistics},
  pages={e1617},
  year={2023},
  publisher={Wiley Online Library}
}

@article{inecik2022multicpa,
  title={MultiCPA: Multimodal Compositional Perturbation Autoencoder},
  author={Inecik, Kemal and Uhlmann, Andreas and Lotfollahi, Mohammad and Theis, Fabian J},
  journal={bioRxiv},
  pages={2022--07},
  year={2022},
  publisher={Cold Spring Harbor Laboratory}
}

@article{janizek2023pause,
  title={PAUSE: principled feature attribution for unsupervised gene expression analysis},
  author={Janizek, Joseph D and Spiro, Anna and Celik, Safiye and Blue, Ben W and Russell, John C and Lee, Ting-I and Kaeberlin, Matt and Lee, Su-In},
  journal={Genome Biology},
  volume={24},
  number={1},
  pages={81},
  year={2023},
  publisher={Springer}
}

@inproceedings{kim2021conditional,
  title={Conditional variational autoencoder with adversarial learning for end-to-end text-to-speech},
  author={Kim, Jaehyeon and Kong, Jungil and Son, Juhee},
  booktitle={International Conference on Machine Learning},
  pages={5530--5540},
  year={2021},
  organization={PMLR}
}

@article{chen2019robust,
  title={Robust attribution regularization},
  author={Chen, Jiefeng and Wu, Xi and Rastogi, Vaibhav and Liang, Yingyu and Jha, Somesh},
  journal={Advances in Neural Information Processing Systems},
  volume={32},
  year={2019}
}

@inproceedings{sundararajan2017axiomatic,
  title={Axiomatic attribution for deep networks},
  author={Sundararajan, Mukund and Taly, Ankur and Yan, Qiqi},
  booktitle={International conference on machine learning},
  pages={3319--3328},
  year={2017},
  organization={PMLR}
}

@article{he2022human,
  title={A human fetal lung cell atlas uncovers proximal-distal gradients of differentiation and key regulators of epithelial fates},
  author={He, Peng and Lim, Kyungtae and Sun, Dawei and Pett, Jan Patrick and Jeng, Quitz and Polanski, Krzysztof and Dong, Ziqi and Bolt, Liam and Richardson, Laura and Mamanova, Lira and others},
  journal={Cell},
  volume={185},
  number={25},
  pages={4841--4860},
  year={2022},
  publisher={Elsevier}
}

@misc{10xdatasets,
  title = {{10x Genomics Single Cell Gene Expression Datasets}},
  year = {2023},
  author = {{10x Genomics}},
  howpublished = {Retrieved from \textnormal{\url{https://support.10xgenomics.com/single-cell-gene-expression/datasets}}},
  note = {Accessed: 2023-05-12}
}

@article{gayoso2022python,
  title={A Python library for probabilistic analysis of single-cell omics data},
  author={Gayoso, Adam and Lopez, Romain and Xing, Galen and Boyeau, Pierre and Valiollah Pour Amiri, Valeh and Hong, Justin and Wu, Katherine and Jayasuriya, Michael and Mehlman, Edouard and Langevin, Maxime and others},
  journal={Nature biotechnology},
  volume={40},
  number={2},
  pages={163--166},
  year={2022},
  publisher={Nature Publishing Group US New York}
}

@article{wolf2018scanpy,
  title={SCANPY: large-scale single-cell gene expression data analysis},
  author={Wolf, F Alexander and Angerer, Philipp and Theis, Fabian J},
  journal={Genome biology},
  volume={19},
  pages={1--5},
  year={2018},
  publisher={Springer}
}

@article{luecken2022benchmarking,
  title={Benchmarking atlas-level data integration in single-cell genomics},
  author={Luecken, Malte D and B{\"u}ttner, Maren and Chaichoompu, Kridsadakorn and Danese, Anna and Interlandi, Marta and M{\"u}ller, Michaela F and Strobl, Daniel C and Zappia, Luke and Dugas, Martin and Colom{\'e}-Tatch{\'e}, Maria and others},
  journal={Nature methods},
  volume={19},
  number={1},
  pages={41--50},
  year={2022},
  publisher={Nature Publishing Group US New York}
}
