## Supplementary material for "scARE: Attribution Regularization for Single Cell Representation Learning": LaTeX: example_paper.tex

%%%%%%%% ICML 202 EXAMPLE LATEX SUBMISSION FILE %%%%%%%%%%%%%%%%%

\documentclass[nohyperref]{article}

\usepackage{lipsum}

% Recommended, but optional, packages for figures and better typesetting:
\usepackage{microtype}
\usepackage{graphicx}
%\usepackage{subfigure}
\usepackage{booktabs} % for professional tables
\usepackage{caption}
\usepackage{subcaption}

% hyperref makes hyperlinks in the resulting PDF.
% If your build breaks (sometimes temporarily if a hyperlink spans a page)
% please comment out the following usepackage line and replace
% \usepackage{icml2023} with \usepackage[nohyperref]{icml2023} above.
\usepackage{hyperref}
\usepackage{url}

% Attempt to make hyperref and algorithmic work together better:
\newcommand{\theHalgorithm}{\arabic{algorithm}}

% Use the following line for the initial version submitted for review:
% \usepackage{icml2023}

% If accepted, instead use the following line for the camera-ready submission:
\usepackage[accepted]{icml2023}

% For theorems and such
\usepackage{amsmath}
\usepackage{amssymb}
\usepackage{mathtools}
\usepackage{amsthm}

% if you use cleveref..
\usepackage[capitalize,noabbrev]{cleveref}

% Redefine the URL link style
\hypersetup{
    colorlinks = true,
    urlcolor = ., % Set the same color as the text
}
\urlstyle{same}


%%%%%%%%%%%%%%%%%%%%%%%%%%%%%%%%
% THEOREMS
%%%%%%%%%%%%%%%%%%%%%%%%%%%%%%%%
\theoremstyle{plain}
\newtheorem{theorem}{Theorem}[section]
\newtheorem{proposition}[theorem]{Proposition}
\newtheorem{lemma}[theorem]{Lemma}
\newtheorem{corollary}[theorem]{Corollary}
\theoremstyle{definition}
\newtheorem{definition}[theorem]{Definition}
\newtheorem{assumption}[theorem]{Assumption}
\theoremstyle{remark}
\newtheorem{remark}[theorem]{Remark}

% Todonotes is useful during development; simply uncomment the next line
%    and comment out the line below the next line to turn off comments
%\usepackage[disable,textsize=tiny]{todonotes}
\usepackage[textsize=tiny]{todonotes}


% The \icmltitle you define below is probably too long as a header.
% Therefore, a short form for the running title is supplied here:
\icmltitlerunning{scARE: Attribution Regularization for Single Cell Representation Learning}
% cellruler

\begin{document}

\twocolumn[
\icmltitle{scARE: Attribution Regularization for Single Cell Representation Learning}


% List of affiliations: The first argument should be a (short)
% identifier you will use later to specify author affiliations
% Academic affiliations should list Department, University, City, Region, Country
% Industry affiliations should list Company, City, Region, Country

% You can specify symbols, otherwise they are numbered in order.
% Ideally, you should not use this facility. Affiliations will be numbered
% in order of appearance and this is the preferred way.
\icmlsetsymbol{equal}{*}

\begin{icmlauthorlist}
\icmlauthor{Kemal Inecik}{icb,weih}
\icmlauthor{Fabian Theis}{icb,matum}
\end{icmlauthorlist}

\icmlaffiliation{icb}{Institute of Computational Biology, Helmholtz Center Munich, Neuherberg, Germany}
\icmlaffiliation{matum}{Department of Mathematics, Technical University of Munich, Garching, Germany}
\icmlaffiliation{weih}{School of Life Sciences Weihenstephan, Technical University of Munich, Freising, Germany}

\icmlcorrespondingauthor{Fabian Theis}{}

% You may provide any keywords that you
% find helpful for describing your paper; these are used to populate
% the "keywords" metadata in the PDF but will not be shown in the document
\icmlkeywords{Machine Learning, ICML, feature attributions, attribution priors, attribution loss, expected gradients, integrated gradients, VAE, cVAE, autoencoder, single-cell}

\vskip 0.3in
]

% this must go after the closing bracket ] following \twocolumn[ ...

% This command actually creates the footnote in the first column
% listing the affiliations and the copyright notice.
% The command takes one argument, which is text to display at the start of the footnote.
% The \icmlEqualContribution command is standard text for equal contribution.
% Remove it (just {}) if you do not need this facility.

\printAffiliationsAndNotice{}  % leave blank if no need to mention equal contribution
%\printAffiliationsAndNotice{\icmlEqualContribution} % otherwise use the standard text.

\begin{abstract}

Single-cell data generation techniques have provided valuable insights into the intricate nature of cellular heterogeneity. However, effectively unraveling subtle variations within a specific gene set of interest, while mitigating the confounding presence of higher-order variability, remains challenging. To address this, we propose scARE, a novel end-to-end generative deep learning model, amplifies model sensitivity to a preselected subset of features while minimizing others. scARE incorporates an auxiliary attribution loss term during model training, which empowers researchers to manipulate the model's behavior robustly and flexibly. In this study, we showcased scARE's applicability in two concrete scenarios: uncovering subclusters associated with the expression patterns of two cellular pathway genes, and its ability to optimize the model training procedure by leveraging time-points metadata, resulting in improved downstream performance.

\end{abstract}

\vspace{-2.0em}
\section{Introduction} \label{sec:introduction}
\vspace{-0.4em}

% The advent of single-cell data generation methods has sparked a paradigm shift, enabling the exploration of the complex heterogeneity present within cell populations, and providing unprecedented insights into the molecular and functional characteristics of individual cells. 
Deep neural networks, particularly autoencoders, are extensively employed in integrating and analyzing single-cell data, demonstrating outstanding performance in tasks such as batch correction, dimension reduction, and perturbation modeling \cite{lopez2018deep, inecik2022multicpa, heumos2023best}. While biologically informed deep learning is an active research area \citep{lotfollahi2023biologically, qoku2023encoding, conard2023spectrum, janizek2023pause}, a method is currently lacking that enables robust and flexible manipulation of an autoencoder model's behavior in order to enhance the influence of a \textit{freely} chosen subset of input features on the latent space or the reconstruction process. In this study, we present scARE, \textit{s}ingle \textit{c}ell \textit{a}ttribution \textit{re}gulatization, an end-to-end generative model designed to amplify the model's sensitivity to a predefined subset of features while concurrently reducing sensitivity to others. It employs a novel training procedure for single-cell data through an auxiliary \textit{attribution loss} term, aimed at effectively unveiling the intricate cellular heterogeneity sought after, while minimizing the potential interference of confounding higher-order variations.

We demonstrate that scARE enables the formation of distinct subclusters for each cell-type in latent space, which differentiate from one another based on the expression patterns associated with the cellular pathway of interest. Enabling unrestricted selection of feature subsets, scARE provides the opportunity to investigate cellular heterogeneity in the expression patterns not only of genes from established biological pathways but also of custom gene lists of interest to researchers. In addition, feature selection based on their association with diseases (such as cancer, diabetes, or viral infections) could potentially facilitate the exploration of diverse cellular states that may be intricately linked to distinct responses and sensitivities to these diseases. 

Moreover, scARE allows users to guide model training by utilizing the metadata of each data point,
%(e.g. selecting different marker genes for each cell-type as the feature subset)
potentially leading to improved performance in downstream analyses. We show that scARE is able to steer model training to segregate clusters populated by cells from different time-points for each cell-type. While yielding outcomes similar to cVAE approaches that utilize additional information regarding cell-type labels during model training \cite{xu2021probabilistic, de2022population}, scARE achieves greater flexibility by employing binary attribution prior vectors instead of relying solely on such labels. Furthermore, we show that scARE adeptly learns a probabilistic representation of the data while concurrently addressing biological and technical factors, achieving performance on par with the baseline model.

\vspace{-0.65em}
\section{Methods} \label{sec:methods}
\vspace{-0.4em}
% it should be 4 pages long. so no text in between here.

\subsection{scARE model architecture}
\vspace{-0.25em}

We propose to incorporate an auxiliary loss term into a conditional variational autoencoder (cVAE) as the base model, where the conditions correspond to batch labels and are provided to both the generative and inference models. cVAEs have been shown to effectively handle the inherent sparsity, noise, and variability in single-cell data while capturing latent representations that facilitate various downstream analyses such as clustering, data imputation, trajectory inference, and label transfer \cite{heumos2023best}. Both generative and inference models are optimized jointly by maximizing the evidence lower bound, ELBO, of the intractable marginal log-likelihood of data $\log p_{\theta}(x|c)$ given in Equation~\ref{eq:elbo}. 

\begin{equation} %
\footnotesize
\setlength{\abovedisplayskip}{-1.6em}
\setlength{\belowdisplayskip}{-0.2em}
\begin{split} \label{eq:elbo}
    \raisetag{1.30\baselineskip}
    \log p_{\theta}(x|c) &\geq \mathbb{E}_{q_{\phi}(z|x,c)} \left[ \log p_{\theta}(x|z,c) - \log \frac{q_{\phi}(z|x,c)}{p_{\theta}(z|c)} \right] \quad 
\end{split}
\end{equation}

Here, $p_{\theta}(z|c)$ denotes a prior distribution of the latent variables $z$ given condition $c$, $p_{\theta}(x|z,c)$ is the likelihood function of a data point $x$, $q_{\phi}(z|x,c)$ is an approximate posterior distribution, and $\theta$ and $\phi$ are respectively generative and inference model parameters. The training loss is defined as the negative ELBO, which can be understood as the sum of the reconstruction loss, $\log p_{\theta}(x|z,c)$, and KL divergence, $\log q_{\phi}(z|x,c)-\log p_{\theta}(z|c)$, where latent $z$ is sampled using the re\-parameterization trick \cite{kingma2015variational, kim2021conditional} from the posterior.

A separate encoder for size factors is implemented, similar to scVI \cite{lopez2018deep}, to enhance the model's performance, interpretability in downstream analyses, and robustness to noise and heterogeneity. The generative model operates on the latent space to reconstruct observed data using a negative binomial loss function (Equation~\ref{eq:loss_r}) specified by mean $\mu$ and inverse dispersion $\theta$. KL divergence was calculated assuming Normal distribution in the latent space (Equation~\ref{eq:loss_kl}), with $\mu_l$ and $\sigma_l$ estimated by the inference model.

\begin{footnotesize}
\setlength{\abovedisplayskip}{-1.5em}
\setlength{\belowdisplayskip}{-0.2em}
\begin{align} % Use the align environment
    % Equation 1
    L_{\mathrm{R}} 
    &= \mathrm{NB}(x; \mu, \theta) \nonumber \\[-0.1em] % Add \nonumber to prevent numbering of intermediate lines
    &= \frac{\Gamma(x+\theta)}{\Gamma(x+1) \Gamma(\theta)}\left(\frac{\theta}{\theta+\mu}\right)^{\theta}\left(\frac{\mu}{\theta+\mu}\right)^{x} \label{eq:loss_r} \\
    % Equation 2
    L_{\mathrm{D}} 
    &= - \mathrm{KL}(\mathcal{N}(\mu_l, \sigma_l) \lVert \mathcal{N}(0, 1)) \label{eq:loss_kl}
\end{align}
\end{footnotesize}

In this study, an auxiliary loss term, attribution loss,  $L_{\mathrm{A}}$, is introduced and incorporated into the base model. The modified model aims to have higher sensitivity to variations in a specific set of input feature values while maintaining its known effectiveness as a cVAE in handling the complex characteristics of single-cell data. The total loss function for the resulting model can be formulated as in Equation~\ref{eq:totalloss}, where $w_A$, $w_D$, and $w_R$ are hyperparameters that dictate the relative contributions of each loss component.

\begin{footnotesize}
\setlength{\abovedisplayskip}{-1.3em}
\setlength{\belowdisplayskip}{-0em}
\begin{align} % Use the align environment
    L_{\text{scARE}} 
    &= L_{\mathrm{R}} w_R + L_{\mathrm{D}} w_D + L_{\mathrm{A}} w_A  \label{eq:totalloss}
\end{align}
\end{footnotesize}

\vspace{-0.5em}
\subsection{Attribution loss}
\vspace{-0.25em}

To compute the attribution loss, $ L_{\mathrm{A}}$, feature attribution methods expected gradients (EG) or integrated gradients (IG) are employed on individual minibatches as the first step. Given a differentiable function $F: \mathbb{R}^n \rightarrow \mathbb{R}$, where $n$ is the number of input features, IG aims to attribute the difference in the function's output between a given reference input $x'$, typically a zero vector, and the input of interest $x$ to the individual features of $x$ \cite{sundararajan2017axiomatic, chen2019robust}. The attribution for the $i_{\mathrm{th}}$ feature is defined in Equation~\ref{eq:ig}, where the integral computes the average gradient along the straight-line path from $x'$ to $x$ in the input space.

\begin{footnotesize}
\setlength{\abovedisplayskip}{-1.6em}
\setlength{\belowdisplayskip}{-0.1em}
\begin{align} % Use the align environment
    % Equation 1
    \text{IG}_{i}(x, x') 
    &= (x_{i} - x') \int_{\alpha=0}^1 \frac{\partial F(x' + \alpha (x - x'))}{\partial x_{i}} \, d\alpha \label{eq:ig}
\end{align}
\end{footnotesize}

EG computes the expected contribution of each feature to the function's output over a distribution of reference inputs \cite{erion2021improving}. Given a distribution $D$ over reference inputs $x'$, the EG attribution for the $i_{\mathrm{th}}$ feature is defined in Equation~\ref{eq:eg}. A simple notation is also in Equation~\ref{eq:eg} introducing noise distribution directly, using a variable $\epsilon$ representing a random perturbation drawn from a distribution $D_{\epsilon}$. In this study, EG was preferred over IG on account of its robustness to reference inputs, ability to handle non-linearities and feature interactions, provision of distribution-aware explanations, and its superior efficiency with expedited model training process \cite{hesse2021fast, erion2021improving}.

\begin{footnotesize}
\setlength{\abovedisplayskip}{-1.0em}
\setlength{\belowdisplayskip}{-0.2em}
\begin{align} % Use the align environment
    % Equation 1
    \text{EG}_{i}(x) 
    &= \mathbb{E}_{x' \sim \text{D}, \alpha \sim U(0,1)} \left[ (x_{i} - x') \frac{\partial F(x' + \alpha (x - x'))}{\partial x_{i}} \right]  \nonumber   \\[-0.05em]
    &= \mathbb{E}_{\epsilon \sim \text{D}_{\epsilon}} \left[\frac{\partial F(x_{i} + \epsilon)}{\partial x_{i}}\right]  \label{eq:eg} 
\end{align}
\end{footnotesize}

The function $F$ used in Equations~\ref{eq:ig} and~\ref{eq:eg} serves to consolidate the inference or generative processes into a single scalar, thereby enabling the regularization of either the entire model or exclusively the encoder during the training phase through attribution loss. While various forms of $F$ have been implemented, including the simplest one being a summation operation, the selection of $F$ remains a model hyperparameter to be tuned. After calculating attributions, the resulting vector undergoes a transformation to ensure non-negativity and scaling to reside within the interval $[0, 1]$. Several re-\-parameterization methods, such as sigmoid transformation or absolute value operation followed by min-max scaling, were experimented in this context. The selection of an effective re-\-parameterization method is a critical hyperparameter, requiring careful tuning due to its direct impact on model performance.

Prior to commencing the training procedure, a subset of input features must be chosen based on the objectives of subsequent downstream analysis. As previously mentioned, there are no predefined regulations or restrictions governing feature selection, granting the user absolute autonomy to make choices aligned with their analysis objectives. scARE requires a binary vector, called ground truth, \textit{GT}, where features to sensitize are assigned $1$ and the remaining features are assigned $0$. Alternatively, features can be selected independently for individual data points, resulting in a binary GT matrix. In this study, we chose genes associated with {\footnotesize\texttt{GO:0043410}} and {\footnotesize\texttt{GO:1901224}} Gene Ontology (GO) terms as GT gene definitions in Section~\ref{subsec:subclustering}, which correspond to positive regulation of MAPK and NF-$\kappa$B pathways respectively. The selection of these specific GO terms was solely intended to demonstrate the proof-of-concept of a biological application. However, it is noteworthy that comparable results were achieved when employing different GO terms (data not presented). On the other hand, in Section~\ref{subsec:segregation}, the features are chosen by their established role as cell-type markers, or their ability to explain the variability across different time-points of each cell-type.

Lastly, the predefined GT vector is compared with the calculated attribution vector for each data point typically through MSE or MAE as in Equation~\ref{eq:msemae}, where $n$ is number of features, $g_i$ and $a_i$ are respectively predefined GT and calculated attribution values. We designed attribution loss to penalize the discrepancy between expected attributions defined by GT vector with actual attribution scores, constraining the model to prioritize the variation in the chosen feature values while de-emphasizing the variation in other features. scARE also allows multiple GTs, where individual attribution losses are computed separately, potentially employing different functions. The total attribution loss is then aggregated, either by simple summation or squared-summation, incorporating a weight term for each attribution loss component. 

\begin{footnotesize}
\setlength{\abovedisplayskip}{-1.5em}
\setlength{\belowdisplayskip}{-0.25em}
\begin{align} \label{eq:msemae}% Use the align environment
    % Equation 1
    \begin{split}
    L_{A_{\text{MSE}}} &= \frac{1}{n} \sum_{i=1}^{n} (g_i - a_i)^2
    \end{split}
    && \hspace{-0.5em}
    % Equation 2
    \begin{split}
    L_{A_{\text{MAE}}} &= \frac{1}{n} \sum_{i=1}^{n} |g_i - a_i|
    \end{split}
\end{align}
\end{footnotesize}

\vspace{-0.6em}
\subsection{Feature-set subcluster founder score}
\vspace{-0.25em}

We devised a novel metric, named feature-set subcluster founder score, \textit{FSSF}, to investigate whether subclusters are formed within a given cell-type based on the expression patterns of GT genes. To calculate this metric, an experimental process is executed wherein the expression values for GT genes are swapped between cells belonging to two distinct subclusters of a cell-type, which have been previously identified by the Leiden algorithm in the latent space. The resulting modified cells, which were unseen during the model training phase, now carry the GT gene expression values from the cells of another latent subcluster. The FSSF score aims to quantify the degree to which these cells map in the boundaries of the subcluster from which the GT genes were sourced, as well as to the subcluster from which all other genes were sourced. The primary objective is to evaluate whether the model perceives all cells within these two subclusters as identical, with the sole differentiating feature being the patterns of GT gene expression. For each pair of Leiden subclusters within a cell-type, the FSSF score is formulated as $(n_g - n_o) / n_t$. Here, $n_g$ and $n_o$ represent the cell count that remapped to the subcluster where the GT genes belong and to the subcluster where all other genes belong, respectively, while $n_t$ represents the total number of cells involved in the exchange. The FSSF score hence is a measure that indicates the relative impact of GT genes versus all other genes on the identity of a cell-type's subclusters. A score close to $1$ implies that GT genes dominate this identity, while a score near $-1$ signifies that all other genes exert a more significant influence.


\vspace{-0.6em}
\subsection{Datasets}
\vspace{-0.25em}

In this study, two scRNA-seq datasets were used for model training and subsequent analyses. The first dataset, named \textit{PBMC}, was provided and processed by \textit{scvi-tools} package \cite{gayoso2022python, 10xdatasets}, resulting in a dataset with $11990$ cells and $3346$ genes. The second dataset, named \textit{He2022} \cite{he2022human}, underwent quality checking and filtering for samples from $15$ and $22$ post-conception weeks, resulting in $8963$ cells, followed by the selection of $2000$ highly variable genes (HVGs) using \textit{scanpy} \cite{wolf2018scanpy}. 

\vspace{-0.65em}
\section{Results} \label{sec:results}
\vspace{-0.4em}

\subsection{Robust subclustering in cell-types associated with expression patterns of predefined gene subsets} \label{subsec:subclustering}
\vspace{-0.25em}

\begin{figure}[!tb]%+
    \begin{subfigure}[t]{1\columnwidth}
        \centering
        \includegraphics[width=0.85\textwidth]{figures/subcluster_1.pdf}
        \vspace{-0.15em}
        \subcaption{}
        \label{fig:subcluster_boxplot}
    \end{subfigure}
    \begin{subfigure}[t]{1\columnwidth}
        \centering
        \includegraphics[width=0.75\textwidth]{figures/subcluster_2.pdf}
        \vspace{-0.1em}
        \subcaption{}
        \label{fig:subcluster_examples}
    \end{subfigure}
    \vspace{-1.00em}
    \caption{
    a) Improved FSSF scores upon inclusion of $L_{\mathrm{A}}$. Each column correspond to FSSF scores for all subcluster pairs in cell-types after training. `Others' represents mean score of 100 repetitions of FSSF calculations using randomly selected feature subsets of the same size as GT genes.
    b) Representative cell-type clusters from \textit{He2022} with high FSSF score for each pair subclusters. 
    } 
    \label{fig:subcluster}
    \vspace{-1.25em}
\end{figure}

In conventional single-cell analysis workflows, the discernibility of cellular dissimilarities with respect to a desired set of gene patterns within the latent space is often impeded by a confounding presence of higher-order variability that obfuscates the finer intricacies being pursued. In this study, we showed that incorporating an auxiliary loss term enables steering the model towards a particular set of genes. This leads to the formation of subclusters that correspond to distinct, yet possibly unexplored, cellular states associated with those genes. The prevalence of such subclusters is demonstrated for both datasets, as illustrated in Figure~\ref{fig:subcluster_boxplot}, where FSSF scores approach $1$ exclusively for GT genes when $L_{\mathrm{A}}$ is incorporated. It is worth noting that each of these subclusters exhibits unique expression patterns for GT genes, as substantiated by autocorrelation matrices (data not presented). 

The capabilities of the scARE model manifest clearly in a diverse range of potential applications. It could, for example, help pinpoint unique expression patterns of homeobox genes linked specifically to distinct time and tissue combinations within a large developmental dataset, deepening our understanding of the intricate dynamics of development. Additionally, ongoing research (data not presented) delves into the complex interplay between these cellular states and specific perturbations. Encouragingly, preliminary findings indicate that certain subclusters exhibit heightened responsiveness to particular perturbations, providing valuable insights into diverse cellular behaviors under varying conditions.

Importantly, the proposed approach maintains cellular homogeneity within individual cell-types, similar to the baseline model without attribution loss (further explained in Section~\ref{subsec:scib}), but it results in the reorganization of cells across different Leiden subclusters of a given cell-type (Figure~\ref{fig:subcluster_examples}). Furthermore, although the relationship between the input and latent space is non-linear, the findings demonstrate that the inclusion of attribution loss leads to a  rearrangement of cells in the latent space such that they display a gradient-like polarization in the total expression of GT genes (Figure~\ref{fig:polarization}).

\begin{figure}[!b]%+
    \vspace{-0.45em}
    \begin{subfigure}[t]{1\columnwidth}
        \centering
        \includegraphics[width=0.85\textwidth]{figures/subcluster_3.pdf} \quad
    \end{subfigure}
    \vspace{-1.05em}
    \caption{
    Representative cell-type clusters from \textit{He2022}, colored for polarization in terms of total MAPK expression.
    } 
    \label{fig:polarization}
    \vspace{-1.25em}
\end{figure}

\vspace{-0.6em}
\subsection{Steering model training with time-points metadata for improved downstream analysis performance} \label{subsec:segregation}
\vspace{-0.25em}

\begin{figure}[!tb]%+
    \begin{subfigure}[t]{1\columnwidth}
        \centering
        \includegraphics[width=0.85\textwidth]{figures/time_1.pdf}
        \vspace{-0.0em}
        \subcaption{}
        \label{fig:time_boxplot}
    \end{subfigure}
    \begin{subfigure}[t]{1\columnwidth}
        \centering
        \vspace{0.40em}
        \includegraphics[width=0.85\textwidth]{figures/time_2.pdf} \quad 
        \vspace{-0.25em}
        \subcaption{}
        \label{fig:time_umap}
    \end{subfigure}
    \vspace{-0.90em}
    \caption{
    a) Improvement for cluster definition upon incorporation of attribution loss in model training.
    b) Representative cell-type clusters from \textit{He2022} with segregated time-point subclusters by scARE training with cell-type specific time-point markers. Yellow and gray correspond to cells from 15 and 22 PCW respectively.
    } 
    \label{fig:time}
    \vspace{-1.2em}
\end{figure}

Achieving optimal segregation among clusters that potentially delineate distinct phenotypes, such as diverse cell-types, time-points (ages), or organs, constitutes an important element in single-cell analysis workflows, including single-cell atlas building. \cite{luecken2022benchmarking, heumos2023best}. In this study, we showed scARE can reveal subtle changes in the transcriptomic landscape of various cell-types during early human development that are otherwise indistinguishable using conventional methods. scARE enables exploration of the developmental lung dataset, \textit{He2022}, with enhanced performance, allowing for improved gene expression visualization and trajectory inference.

The top 40 genes that account for the variability between two time-points (post-conception week, PCW, 15 and 22) were calculated using mutual information scores for scARE model training. This calculation was performed for all cells, resulting in a GT vector, as well as for each cell-type independently, resulting in a GT matrix. The degree of overlap between time-point subclusters for each cell-type was observed to decrease substantially, leading to a more distinct delineation of cluster definitions (Figure~\ref{fig:time}). The improvement was observed when choosing cell-type specific GTs, which is anticipated given that the genes demarcating separate temporal instances of a cell-type are largely subject to the cell-type itself. Moreover, the initial findings (data not presented) on the HLCA dataset \cite{sikkema2023integrated} suggest that incorporating an attribution loss into the model training, using a GT matrix based on cell-type markers, enhances some model performance metrics such as improved label transfer accuracy and lower prediction uncertainty. 

\vspace{-0.7em}
\subsection{Comparable performance to the baseline model} \label{subsec:scib}
\vspace{-0.3em}

\begin{figure}[!b]%+
    \vspace{-0.55em}
    \begin{subfigure}[t]{1\columnwidth}
        \centering
        \includegraphics[width=0.9\textwidth]{figures/scib_1.pdf}
    \end{subfigure}
    \vspace{-2.2em}
    \caption{
    Data integration performance for batch correction and biological variation conservation, obtained by \textit{scib} \cite{luecken2022benchmarking}. `Unintegrated' is based on the top $50$ PCA components of the input space, while `Only NF-$\kappa$B' is based on the latent space of the baseline model using only these genes.
    } 
    \label{fig:scib}
    \vspace{-1.5em}
\end{figure}

In an ideal scenario, scARE should maintain baseline performance, minimizing any significant compromise, while concurrently delivering substantial enhancements in the interpretability and explication of the model's predictive outcomes. The incorporation of attribution loss was found to have no significant detrimental impact on the effectiveness of data integration, specifically in relation to batch removal and preservation of biological variation (Figure~\ref{fig:scib}).

\vspace{-0.65em}
\section{Discussion} \label{sec:discussion}
\vspace{-0.4em}

The scARE model was shown to effectively improve its own responsiveness to variations in a designated set of feature values while upholding its acknowledged effectiveness as a cVAE for single-cell data. Its practical applications were highlighted in two concrete scenarios: identification of subclusters correlated with gene expression patterns of the predefined gene set, as well as refining the separation of clusters that are presumed to belong distinct groups. The incorporation of attribution loss into cVAE model training, as pioneered by scARE, is expected to aid future investigations into the integration of interpretability and generative modeling techniques.

As part of our future objectives, we aim to support our analysis by showcasing scARE on a broader range of datasets and exploring diverse feature subsets. Through these efforts, we seek to gain deeper insights into the variations among different subclusters and potentially correlate them with pertinent gene programs from the biological literature. Exploring the inclusion of multiple ground truths (e.g. GTs from NF-$\kappa$B and apoptosis pathways simultaneously) would introduce captivating dimensions to the analysis. While our existing implementation enables training with a combined loss, a method still needs to be devised to maintain their disjointness or orthogonality. Furthermore, the use cases of scARE are envisioned to be extended to select an optimal set of probed genes in targeted spatial transcriptomics \cite{kuemmerle2022probe} through the utilization of a zero GT vector and a loss function penalizing larger deviations, thereby imposing a constraint to employ the minimum number of features for reconstruction. As current implementation lacks multimodal functionality, a further objective is to extend scARE to incorporate multimodal single-cell datasets \cite{teichmann2020method}.

\vspace{-0.65em}
\section{Acknowledgement} \label{sec:acknowledgement}
\vspace{-0.4em}

Sincere gratitude is extended to the reviewers for their generous contribution of time and expertise in meticulously evaluating this work. Their insightful feedback and constructive suggestions have significantly elevated the caliber of this paper. Special thanks are also extended to Lisa Sikkema, Louis Kuemmerle, Adil Meric, Ignacio Ibarra, Ekin Gokce Cicek, and Soroor Hediyeh-zadeh for their invaluable feedback during the writing process. Their meticulous attention to detail, thoughtful insights, and astute feedback have played an instrumental role in refining the coherence and effectiveness of this endeavor.

\vspace{-0.65em}
\bibliography{example_paper}
\bibliographystyle{icml2023}

\end{document}


% This document was modified from the file originally made available by
% Pat Langley and Andrea Danyluk for ICML-2K. This version was created
% by Iain Murray in 2018, and modified by Alexandre Bouchard in
% 2019 and 2021 and by Csaba Szepesvari, Gang Niu and Sivan Sabato in 2023. 
% Previous contributors include Dan Roy, Lise Getoor and Tobias
% Scheffer, which was slightly modified from the 2010 version by
% Thorsten Joachims & Johannes Fuernkranz, slightly modified from the
% 2009 version by Kiri Wagstaff and Sam Roweis's 2008 version, which is
% slightly modified from Prasad Tadepalli's 2007 version which is a
% lightly changed version of the previous year's version by Andrew
% Moore, which was in turn edited from those of Kristian Kersting and
% Codrina Lauth. Alex Smola contributed to the algorithmic style files.


%%%%%%%%%%%%%%%%%%%%%%%%%%%%%%%%%%%%%%%%
%%%%%%%%%%%%%%%%%%%%%%%%%%%%%%%%%%%%%%%%


% \vspace{-0.6em}
% \subsection{Datasets}
% \vspace{-0.25em}
% % The introduction of attribution loss often poses obstacles in gradient propagation and causes the model to depend heavily on a limited number of influential features or dimensions. To address these problems, the AdamW optimizer and weight decay are employed in scARE, which stabilize the training process and promote controlled gradient updates and smoother convergence. Additionally, it has been observed that introducing a warm-up period, during which the attribution loss or ELBO component is temporarily excluded, has been experienced to be advantageous for model training in certain dataset and GT combinations. 
% % It is worth mentioning that scARE is developed using the \textit{scvi-tools} framework \cite{gayoso2022python}, ensuring compatibility with existing workflows while extending the applicability of attribution loss to established models like totalVI \cite{gayoso2021joint} and peakVI \cite{ashuach2022peakvi}.
