## Supplementary figures and images for "scARE: Attribution Regularization for Single Cell Representation Learning"

### scib_1.pdf

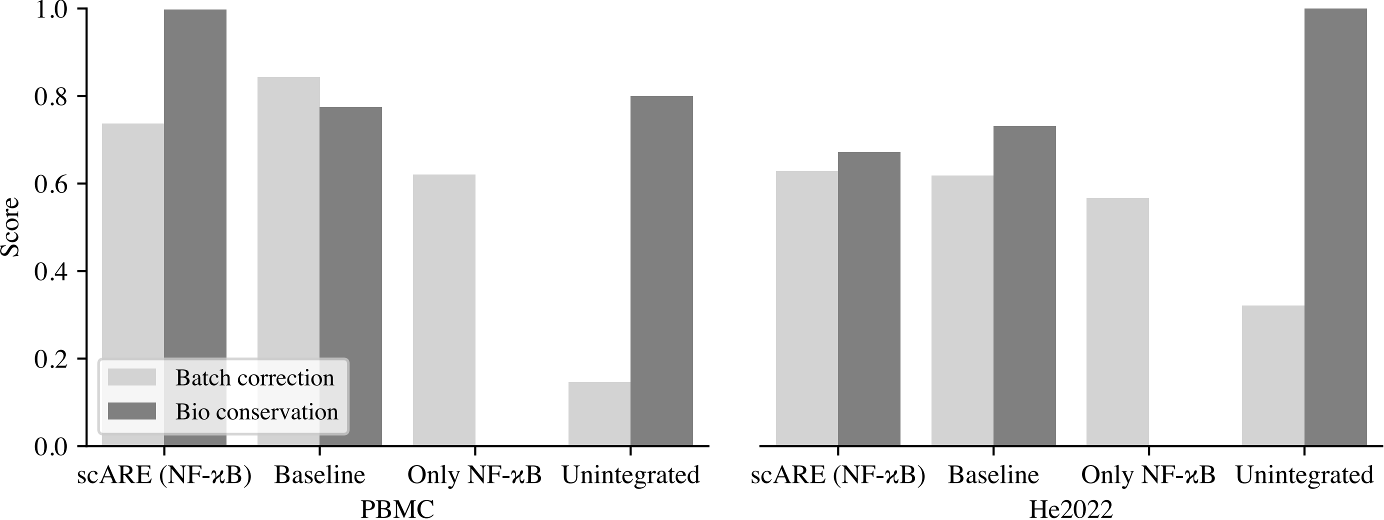

### subcluster_1.pdf

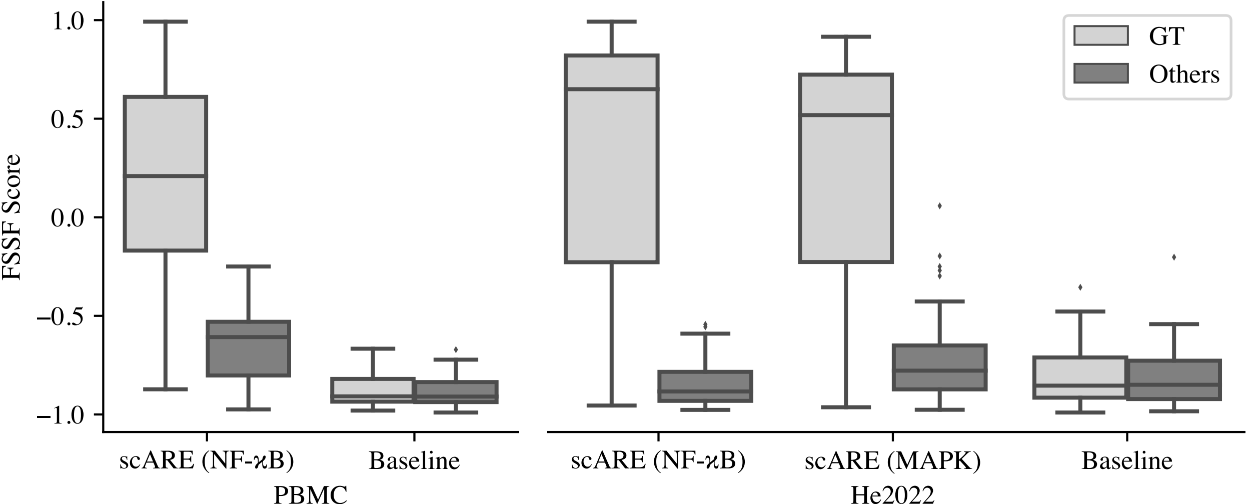

### subcluster_2.pdf

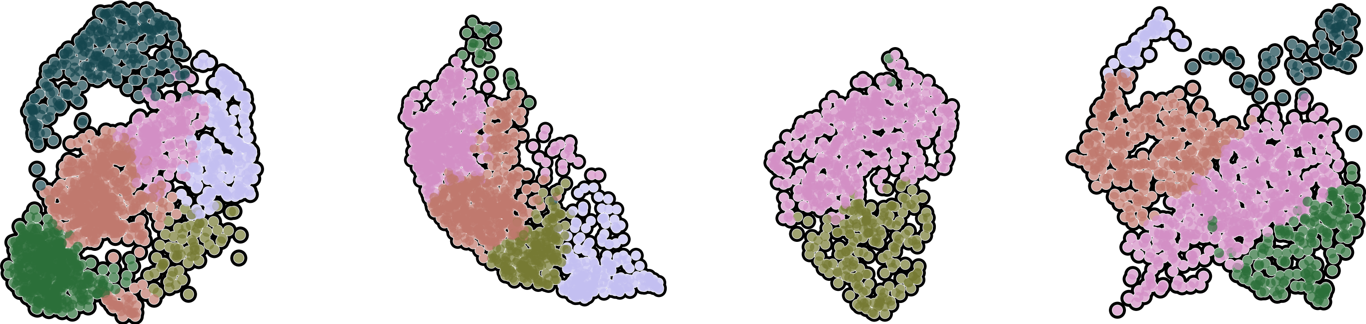

### subcluster_3.pdf

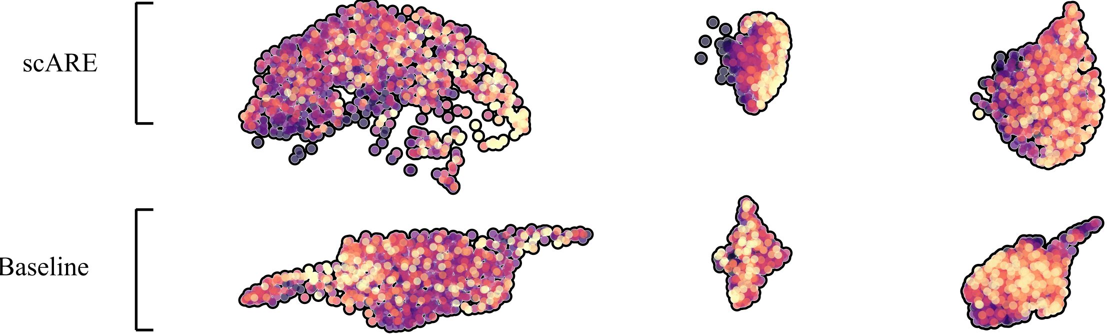

### time_1.pdf

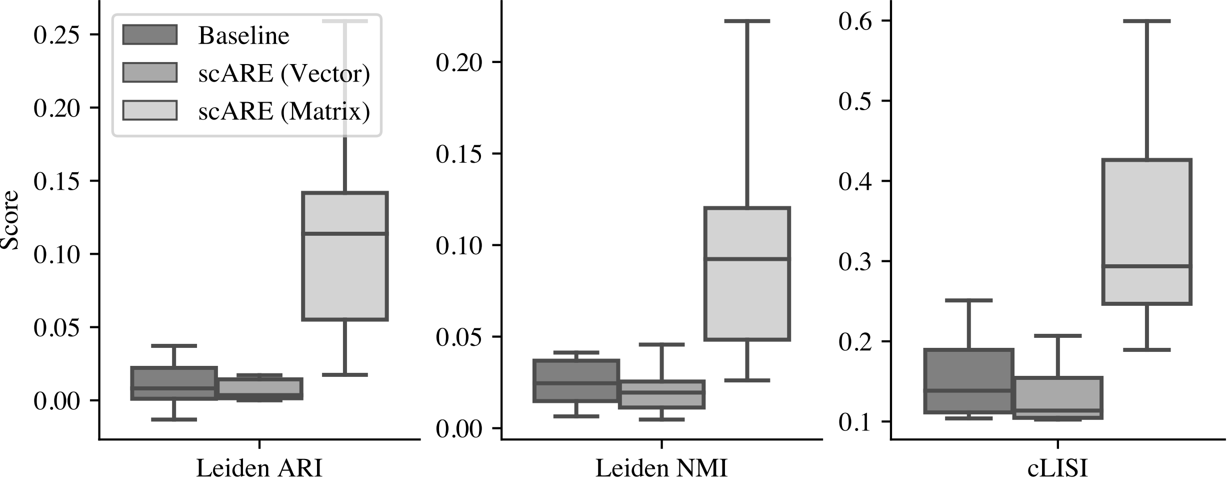

### time_2.pdf

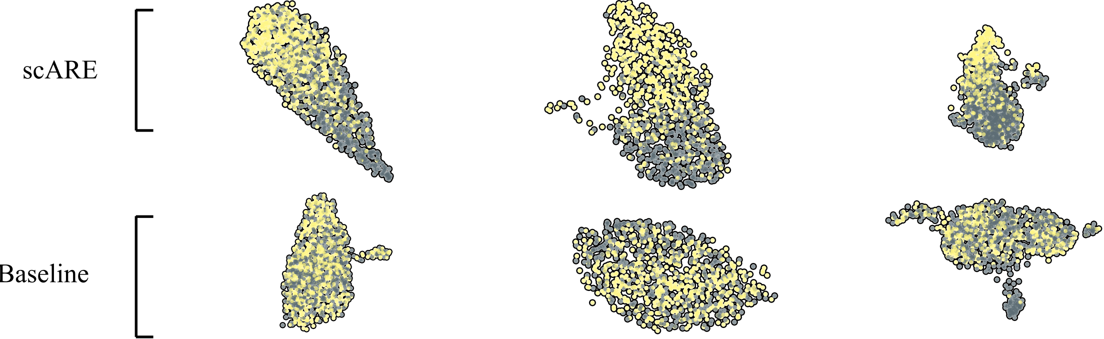
